# Maternal and fetal genetic effects on birth weight and their relevance to cardio-metabolic risk factors

**DOI:** 10.1101/442756

**Authors:** Nicole M Warrington, Robin N Beaumont, Momoko Horikoshi, Felix R Day, Øyvind Helgeland, Charles Laurin, Jonas Bacelis, Shouneng Peng, Ke Hao, Bjarke Feenstra, Andrew R Wood, Anubha Mahajan, Jessica Tyrrell, Neil R Robertson, N William Rayner, Zhen Qiao, Gunn-Helen Øiseth Moen, Marc Vaudel, Carmen J Marsit, Jia Chen, Michael Nodzenski, Theresia M Schnurr, Mohammad H Zafarmand, Jonathan P Bradfield, Niels Grarup, Marjolein N Kooijman, Ruifang Li-Gao, Frank Geller, Tarunveer S Ahluwalia, Lavinia Paternoster, Rico Rueedi, Ville Huikari, Jouke-Jan Hottenga, Leo-Pekka Lyytikäinen, Alana Cavadino, Sarah Metrustry, Diana L Cousminer, Ying Wu, Elisabeth Thiering, Carol A Wang, Christian T Have, Natalia Vilor-Tejedor, Peter K Joshi, Jodie N Painter, Ioanna Ntalla, Ronny Myhre, Niina Pitkänen, Elisabeth M van Leeuwen, Raimo Joro, Vasiliki Lagou, Rebecca C Richmond, Ana Espinosa, Sheila J Barton, Hazel M Inskip, John W Holloway, Loreto Santa-Marina, Xavier Estivill, Wei Ang, Julie A Marsh, Christoph Reichetzeder, Letizia Marullo, Berthold Hocher, Kathryn L Lunetta, Joanne M Murabito, Caroline L Relton, Manolis Kogevinas, Leda Chatzi, Catherine Allard, Luigi Bouchard, Marie-France Hivert, Ge Zhang, Louis J Muglia, Jani Heikkinen, Early Growth Genetics (EGG) Consortium, Camilla S Morgen, Antoine HC van Kampen, Barbera DC van Schaik, Frank D Mentch, Claudia Langenberg, Jian’an Luan, Robert A Scott, Jing Hua Zhao, Gibran Hemani, Susan M Ring, Amanda J Bennett, Kyle J Gaulton, Juan Fernandez-Tajes, Natalie R van Zuydam, Carolina Medina-Gomez, Hugoline G de Haan, Frits R Rosendaal, Zoltán Kutalik, Pedro Marques-Vidal, Shikta Das, Gonneke Willemsen, Hamdi Mbarek, Martina Müller-Nurasyid, Marie Standl, Emil V R Appel, Cilius E Fonvig, Caecilie Trier, Catharina EM van Beijsterveldt, Mario Murcia, Mariona Bustamante, Sílvia Bonas-Guarch, David M Hougaard, Josep M Mercader, Allan Linneberg, Katharina E Schraut, Penelope A Lind, Sarah E Medland, Beverley M Shields, Bridget A Knight, Jin-Fang Chai, Kalliope Panoutsopoulou, Meike Bartels, Friman Sánchez, Jacob Stokholm, David Torrents, Rebecca K Vinding, Sara M Willems, Mustafa Atalay, Bo L Chawes, Peter Kovacs, Inga Prokopenko, Marcus A Tuke, Hanieh Yaghootkar, Katherine S Ruth, Samuel E Jones, Po-Ru Loh, Anna Murray, Michael N Weedon, Anke Tönjes, Michael Stumvoll, Kim F Michaelsen, Aino-Maija Eloranta, Timo A Lakka, Cornelia M van Duijn, Wieland Kiess, Antje Körner, Harri Niinikoski, Katja Pahkala, Olli T Raitakari, Bo Jacobsson, Eleftheria Zeggini, George V Dedoussis, Yik-Ying Teo, Seang-Mei Saw, Grant W Montgomery, Harry Campbell, James F Wilson, Tanja GM Vrijkotte, Martine Vrijheid, Eco JCN de Geus, M Geoffrey Hayes, Haja N Kadarmideen, Jens-Christian Holm, Lawrence J Beilin, Craig E Pennell, Joachim Heinrich, Linda S Adair, Judith B Borja, Karen L Mohlke, Johan G Eriksson, Elisabeth E Widén, Andrew T Hattersley, Tim D Spector, Mika Kähönen, Jorma S Viikari, Terho Lehtimäki, Dorret I Boomsma, Sylvain Sebert, Peter Vollenweider, Thorkild IA Sørensen, Hans Bisgaard, Klaus Bønnelykke, Jeffrey C Murray, Mads Melbye, Ellen A Nohr, Dennis O Mook-Kanamori, Fernando Rivadeneira, Albert Hofman, Janine F Felix, Vincent WV Jaddoe, Torben Hansen, Charlotta Pisinger, Allan A Vaag, Oluf Pedersen, André G Uitterlinden, Marjo-Riitta Järvelin, Christine Power, Elina Hyppönen, Denise M Scholtens, William L Lowe, George Davey Smith, Nicholas J Timpson, Andrew P Morris, Nicholas J Wareham, Hakon Hakonarson, Struan FA Grant, Timothy M Frayling, Debbie A Lawlor, Pål R Njølstad, Stefan Johansson, Ken K Ong, Mark I McCarthy, John RB Perry, David M Evans, Rachel M Freathy

## Abstract

Birth weight (BW) variation is influenced by fetal and maternal genetic and non-genetic factors, and has been reproducibly associated with future cardio-metabolic health outcomes. These associations have been proposed to reflect the lifelong consequences of an adverse intrauterine environment. In earlier work, we demonstrated that much of the negative correlation between BW and adult cardio-metabolic traits could instead be attributable to shared genetic effects. However, that work and other previous studies did not systematically distinguish the direct effects of an individual’s own genotype on BW and subsequent disease risk from indirect effects of their mother’s correlated genotype, mediated by the intrauterine environment. Here, we describe expanded genome-wide association analyses of own BW (n=321,223) and offspring BW (n=230,069 mothers), which identified 278 independent association signals influencing BW (214 novel). We used structural equation modelling to decompose the contributions of direct fetal and indirect maternal genetic influences on BW, implicating fetal- and maternal-specific mechanisms. We used Mendelian randomization to explore the causal relationships between factors influencing BW through fetal or maternal routes, for example, glycemic traits and blood pressure. Direct fetal genotype effects dominate the shared genetic contribution to the association between lower BW and higher type 2 diabetes risk, whereas the relationship between lower BW and higher later blood pressure (BP) is driven by a combination of indirect maternal and direct fetal genetic effects: indirect effects of maternal BP-raising genotypes act to reduce offspring BW, but only direct fetal genotype effects (once inherited) increase the offspring’s later BP. Instrumental variable analysis using maternal BW-lowering genotypes to proxy for an adverse intrauterine environment provided no evidence that it causally raises offspring BP. In successfully separating fetal from maternal genetic effects, this work represents an important advance in genetic studies of perinatal outcomes, and shows that the association between lower BW and higher adult BP is attributable to genetic effects, and not to intrauterine programming.

---

Birth weight (BW) is an important predictor of newborn and infant survival, a key indicator of pregnancy outcomes for mothers as well as for offspring, and is observationally associated with future risk of adult cardio-metabolic diseases in the offspring.

Observational associations between lower BW and later cardio-metabolic diseases are often assumed to reflect adaptations made by a developing fetus in response to an adverse intrauterine environment, such as maternal malnutrition. This concept has been termed the Developmental Origins of Health and Disease (DOHaD) hypothesis^1^. Support of the DOHaD hypothesis is primarily from animal models (reviewed in ^2^). Observational studies of famine-exposed populations support prenatal programming in relation to body size and diabetes, but not other cardio-metabolic health measures (reviewed in ^3^). However, DOHaD cannot provide a complete explanation for the relationship between lower BW and increased risk of cardio-metabolic disease. Other likely contributing factors are (i) environmental confounding, leading to phenotypic associations across the life-course^4^, and (ii) shared genetic effects operating at the population level, as demonstrated in our recent work showing overlap between genetic variants influencing BW and adult cardio-metabolic diseases^5^. Genetic associations between BW and later cardio-metabolic diseases may arise from the direct effects of the same inherited genetic variants at different stages of the life-course^6^. However, consideration of an individual’s own genotype in isolation cannot exclude potential confounding by any indirect effects of the correlated maternal genotype (r≈0.5) on the intrauterine, and possibly postnatal, environment. Evidence for maternal indirect effects on BW and later offspring disease risk could indicate the role of the intrauterine environment in later-life disease etiology.

To date, 65 genomic loci have been associated with BW in genome-wide association studies (GWAS), implicating biological pathways that may underlie observational associations with adult disease^5, 7-9^. However, most of these studies did not distinguish between maternal and fetal genetic influences on BW. Evidence from monogenic human models^10^ and variance components analyses^11^ demonstrate that BW is influenced both by genotypes inherited by the fetus and by maternal genotypes that influence the intrauterine environment. To date, GWAS of own BW^5^ and offspring BW^7^, which have focused on the fetal and maternal genome respectively, have produced overlapping signals due to the correlation between maternal and fetal genotypes. Identified BW variants might have (i) a direct fetal effect only, (ii) an indirect maternal effect only, or (iii) some combination of the two. Performing separate GWAS analyses of own or offspring BW precludes full resolution of the origin of the identified genetic effects. For example, some association signals identified in a GWAS of own BW may in fact be the exclusive consequence of strong indirect maternal effects (and vice versa).

To address these issues, we performed greatly-expanded GWAS of own BW (n=321,223) and offspring BW (n=230,069 mothers) using data from the EGG Consortium and the UK Biobank (2017 release). We applied a statistical method that we recently developed, which utilises structural equation modelling (SEM), to partition genetic effects on BW into maternal and fetal components at genome-wide significant loci^7,12^. We then extended the method to estimate maternal- and fetal-specific genetic effects across the genome in a computationally efficient manner, and used the results for downstream analyses. Our ability to resolve maternal and fetal genetic contributions provides substantial insights into the underlying biological regulation of BW and into the origins of observational relationships with type 2 diabetes (T2D) and blood pressure (BP).

## RESULTS

### Meta-analyses of fetal and maternal GWAS

We conducted GWAS meta-analyses of own (fetal) genetic variants on own BW (**Supplementary Figure 1, Supplementary Tables 1** and **2**) and maternal genetic variants on offspring BW (**Supplementary Figure 2, Supplementary Tables 3** and **4**) in individuals of European ancestry. We then performed approximate conditional and joint multiple-SNP analysis (COJO^13^) and a trans-ethnic meta-analysis to identify further independent SNPs (**Methods**). The GWAS meta-analysis of own BW (N=321,223) identified 211 independent single nucleotide polymorphisms (SNPs) at genome-wide significance (P<5×10^−8^) (**Supplementary Figures 3, 4, 5a, Supplementary Table 5,** and **Methods**). The GWAS meta-analysis of offspring BW (N=230,069 mothers) identified 105 independently associated SNPs (P<5×10^−8^; **Supplementary Figures 3, 4, 5b, Supplementary Table 5,** and **Methods**). When we applied a more stringent significance threshold that accounts for the large number of low frequency SNPs imputed in the UK Biobank and EGG studies (P<6.6×10^−9^; see Kemp et al. 14 for details of the derivation of this threshold), 147 of the 211 SNPs from the GWAS meta-analysis of own BW and 72 of the 105 SNPs from the GWAS meta-analysis of offspring BW remained significant (**Supplementary Table 5**).

SNPs at 52 genome-wide significant loci (within 500Kb) were identified in the GWAS of both own BW and offspring BW. Of these, 11 loci had the same lead SNP and a further 31 loci had fetal and maternal lead SNPs correlated with *r^2^* ≥ 0.1. Colocalization analysis indicated 27/31 of these maternal and fetal lead SNP pairs were likely tagging the same BW signal (posterior probability > 0.5). Therefore, we identified a total of 278 independent association signals, represented by 305 SNPs (**Supplementary Figure 4** and **Supplementary Table 5**). Of the 305 genome-wide significant SNPs, 238 were novel representing 214 independent association signals, four of the identified SNPs are rare (minor allele frequency (MAF)<1%) and 21 are low-frequency (1%≤MAF<5%). Three of the rare variants (*YKT6/GCK*, *ACVR1C* and *MIR146B*) alter BW by more than double the effect (>100g per allele) of the first common variants identified^9^. In the independent Norwegian MoBa-HARVEST study (N=13,934 mother-offspring duos), the variance in BW explained by fetal genetic variation was larger than that explained either by maternal genetic variation or the covariance between the two. The fetal genotype at the genome-wide significant SNPs explained 7% of the variance in BW, whereas the maternal genotype explained 3% and the covariance explained −0.5% (in total, the genome-wide significant SNPs explained 9% of the variance in BW, calculated as the sum of variances explained by the fetal genotype, maternal genotype, plus twice the covariance). Maternal genome-wide complex trait analysis (M-GCTA^11^), which estimates SNP heritability and partitions this quantity into maternal and fetal components, estimated that a total of 39.8% of the variance in BW could be explained by tagged fetal genetic variation (28.5%), tagged maternal genetic variation (7.6%) and twice the covariance between the two (3.7%)..

We integrated data from several sources to highlight possible causal genes underlying the identified associations, including gene-level expression data across 43 tissues (from GTEx v6p^15^), placental expression quantitative trait loci (eQTL^16^), topologically associating domains (TADs) identified in human embryonic stem cells^17,18^ and non-synonymous SNPs (see **Supplementary Table 5** and **Methods**). Several genes were highlighted by multiple approaches; however, further functional studies are required to confirm causality.

### Structural equation model to estimate maternal and fetal effects

We next partitioned the 305 genome-wide significant SNPs into five categories based on their maternal and/or fetal genetic contributions to BW. To achieve this, we used structural equation modelling (SEM) that accounts for the correlation between fetal and maternal genotypes and therefore potential confounding of the maternal and fetal effects on each other^12^. Briefly, the model uses the self-reported BW data from the individuals in the UK Biobank, along with the BW data of the first offspring of the UK Biobank women. We model both grand-maternal and offspring genotypes (which were absent in the UK Biobank) as latent factors, in addition to the genotype data of the UK Biobank individual (**Supplementary Figure 17**). The model is robust to missing data and measurement error, so we could include individuals from the UK Biobank who have reported either their own (including women and men) or their offspring’s BW (women only), but not both. Likewise, we can also include summary statistics from the EGG consortium to improve the estimation of the maternal and fetal effects (see **Methods** for full details). The model provides unbiased estimates of the maternal and fetal genetic effects on BW. We analysed 257,734 individuals of European descent from the UK Biobank (85,518 women with their own and their offspring’s BW, 98,235 men or women with their own BW, and 73,981 women with only their offspring’s BW) and incorporated the summary statistics from the EGG Consortium European meta-analysis of own BW (N=80,745) and offspring BW (N=19,861; **Figure 1**, **Supplementary Figures 4, 6** and **Supplementary Table 5**). Using the confidence intervals around the SEM-adjusted maternal and fetal effect estimates, we identified 83 SNPs with fetal-only effects, 45 SNPs with maternal-only effects, 36 SNPs with directionally-concordant fetal and maternal effects, and 24 SNPs with directionally-opposing fetal and maternal effects (**Supplementary Figure 7**). For example, rs10830963 at *MTNR1B* was identified in both the own BW (P=2.8×10^−11^) and offspring BW (P=9.1×10^−39^) GWAS, but the SEM analysis revealed that its effect on BW was exclusively maternal (P_SEMfetal_=0.7, P_SEMmaternal_=4.6×10^−19^). Conversely, rs28457693 at *PTCH1/FANCC* (own BW GWAS: P=9.9×10^−26^; offspring BW GWAS: P=3.7×10^−9^) showed evidence of a fetal effect only (P_SEMfetal_=1.7×10^−9^, P_SEMmaternal_=0.2). SNP rs560887 at *G6PC2* was identified only in the GWAS of offspring BW (P=1.2×10^−14^), but was found to have directionally-opposing maternal and fetal effects on BW (P_SEMfetal_=2.8×10^−8^, P_SEMmaternal_=5.4×10^−14^). At present, these categories are suggestive as the current sample size has insufficient statistical power to detect small genetic effects, particularly maternal effects. There were 117 SNPs that were unclassified, and some of the SNPs that were classified as fetal only, for example, may have had a small maternal effect that was undetected with the current sample size. Asymptotic power calculations showed that with the current sample size we had 80% power to detect fetal (maternal) effects that explained 0.006% (0.008%) of the variance in BW (α=0.05). However, there was strong consistency with traditional conditional linear regression modelling in N=18,873 mother-offspring pairs (**Supplementary Table 6** and **Methods**), and overall, the method gave a clear indication as to which genetic associations are driven by the maternal or fetal genomes, respectively.

**Figure 1:**
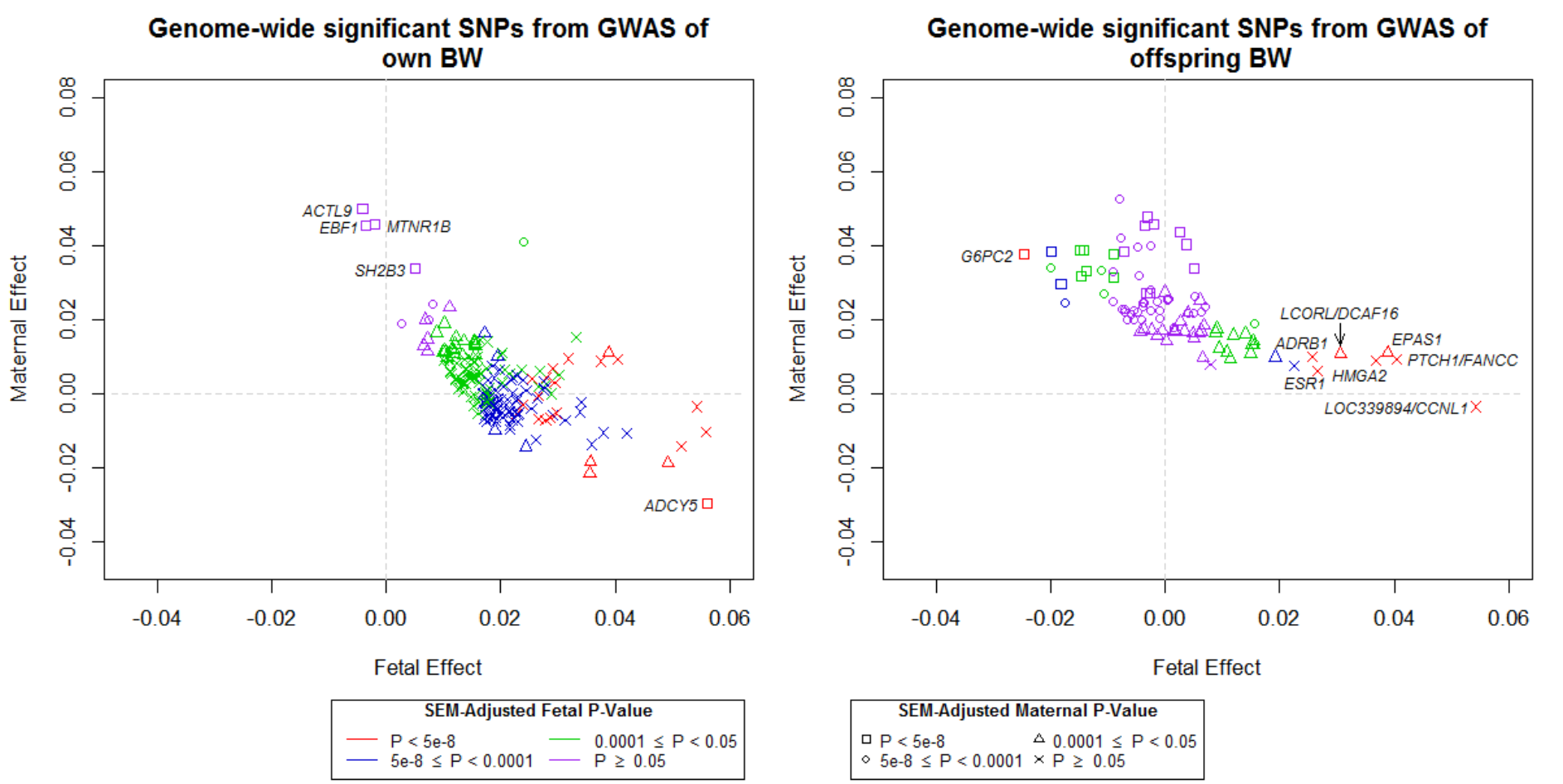
Structural equation modelling (SEM)-adjusted fetal and maternal effects for the 289 genome-wide significant SNPs that were identified in the GWAS of either own birth weight (BW; left panel) or offspring BW (right panel) with minor allele frequency greater than 5%. The colour of each point indicates the SEM-adjusted fetal effect on own BW association P-value and the shape of each point indicates the SEM-adjusted maternal effect on offspring BW association P-value. SNPs which are labelled with the name of the closest gene are those which were identified in the GWAS of own BW but whose effects are mediated through the maternal genome (left panel) and SNPs that were identified in the GWAS of offspring BW but whose effects are mediated through the fetal genome (right panel). SNPs are aligned to the BW increasing allele from the GWAS.

To extend the estimates of adjusted maternal and fetal effects genome-wide, we developed a weighted linear model (WLM) that yields a good approximation to the SEM (see **Methods**), with equivalent estimates for the 305 genome-wide significant SNPs (**Supplementary Figure 8**). This was necessary because the SEM is too computationally intensive to fit to a large number of SNPs across the genome. The resulting adjusted fetal and maternal genotype effect estimates on BW from the WLM are hereafter referred to as WLM-adjusted estimates. Using linkage disequilibrium (LD) score regression^19^, we observed that the genetic correlation between the WLM-adjusted maternal and fetal effects (r_g_=0.10, P=0.12) was substantially lower than that between the unadjusted effects from the original GWAS (r_g_=0.82, P<0.01), indicating that the WLM largely accounts for the underlying correlation between fetal and maternal genotypes. No additional novel loci were identified in the WLM-adjusted analyses. We used these WLM-adjusted estimates in downstream analyses to identify fetal-specific and maternal-specific mechanisms that regulate BW and to investigate the genetic links between BW and adult traits.

### Maternal- and fetal-specific tissues and mechanisms underlying BW regulation

Using the WLM-adjusted estimates, we observed differences in enrichment between the maternal and fetal profiles of gene expression across tissues, and of regulatory pathways. Tests of global enrichment of BW SNP associations across tissues sampled from the GTEx project^15^ using LD-SEG^20^, indicated that the only tissues reaching significance after Bonferroni correction was enrichment for maternal-specific SNP associations for genes expressed in connective/bone tissues (**Supplementary Figure 9**). Integration of epigenetic signatures defined by the Roadmap Epigenomics project highlighted, after Bonferroni correction, a significant enrichment of maternal-specific effects in the ovary for histone modification marks (H3K4me1) and regions of open chromatin (**Supplementary Table 7**); no significant enrichment was detected for other signatures. Gene-set enrichment analysis using WLM-adjusted effect estimates also implicated different gene sets having fetal-specific influences on BW (**Supplementary Table 8**) to those having maternal-specific influences (**Supplementary Table 9)**.

A major determinant of BW is the duration of gestation. We performed LD score regression analysis^19^ to investigate the genetic correlation between published maternal genotype effects on gestational duration^21^ and the WLM-adjusted BW effects. (To date, there is no published GWAS of fetal genotype and gestational duration.) We found a substantial genetic correlation with the WLM-adjusted maternal effects on offspring BW (r_g_=0.63; P=2.1×10^−5^; **Supplementary Table 10**; see **Methods**), but not with the WLM-adjusted fetal effects on own BW (r_g_=-0.10, P=0.35). Gestational duration was unavailable for >85% of individuals in the GWAS analyses, due to the large UK Biobank sample without gestational duration, so it is possible that some identified association signals influence BW primarily by altering the timing of delivery. We looked up the 305 genome-wide significant BW-associated SNPs in the published maternal GWAS of gestational duration^21^ (**Supplementary Table 11**) and followed up 6 SNPs in 13,206 mother-child pairs (P<1.6×10^−4^ with gestational duration, corrected for 305 tests, **Methods**). Meta-analyzing the results from the mother-child pairs with summary data from 23andMe^21^ strengthened associations with gestational duration at four of the six loci (*EBF1, AGTR2, ZBTB38* and *KCNAB1*; **Supplementary Table 12**). The precise causal relationship between fetal growth and gestational duration at these loci requires further investigation, however, the majority of associations with BW do not appear to be driven by associations with gestational duration.

### Maternal- and fetal-specific genetic correlations between BW and adult traits

The 305 genome-wide significant BW-associated SNPs were collectively associated with a wide variety of other phenotypes in previously-published GWAS and in the UK Biobank (**Supplementary Table 13**; see **Methods**). At the genome-wide level, we previously reported genetic correlations between own BW and several adult cardio-metabolic disease traits^5^, for example systolic blood pressure (SBP; r_g_=-0.22, P=5.5×10^−13^), but at that time were unable to distinguish the direct fetal genotype contribution from the indirect contribution of maternal genotype. To understand these distinct contributions, we calculated genetic correlations using LD score regression^19^ between WLM-adjusted fetal and maternal SNP effect estimates and GWAS estimates for a large range of health-related traits (**Figure 2, Supplementary Table 10** and **Methods)**. For many traits, for example adult height, the WLM-adjusted fetal effect on own BW (r_g_=0.28, P=8.1×10^−16^) showed a similar genetic correlation to the WLM-adjusted maternal effect on offspring BW (r_g_=0.29, P=5.1×10^−16^). However, for others, we observed differing fetal-specific and maternal-specific genetic correlations. For example, for several glycemic traits (T2D, 2-hour glucose, fasting glucose, fasting insulin), there were directionally-opposite fetal (own BW) and maternal (offspring BW) genetic correlations. Moreover, the genetic correlations with glycemic traits that were estimated using the WLM-adjusted effects were substantially larger than those estimated using the unadjusted effects, demonstrating the importance of accounting for the maternal-fetal genotype correlation (e.g. fasting glucose: WLM-adjusted fetal effect on own BW r_g_=-0.25, P=8.2×10^−6^; unadjusted fetal effect on own BW r_g_=-0.11, P=0.005; WLM-adjusted maternal effect on offspring BW r_g_=0.20, P=0.003; unadjusted maternal effect on offspring BW r_g_=0.08, P=0.09). Cardiovascular traits showed directionally consistent WLM-adjusted maternal and WLM-adjusted fetal genetic correlations, but with different strengths. For example, the genetic correlation between SBP and WLM-adjusted maternal effects on offspring BW (r_g_=-0.23, P=9.2×10^−10^) was stronger than that between SBP and WLM-adjusted fetal effects on own BW (r_g_=-0.14, P=9.8×10^−5^).

**Figure 2:**
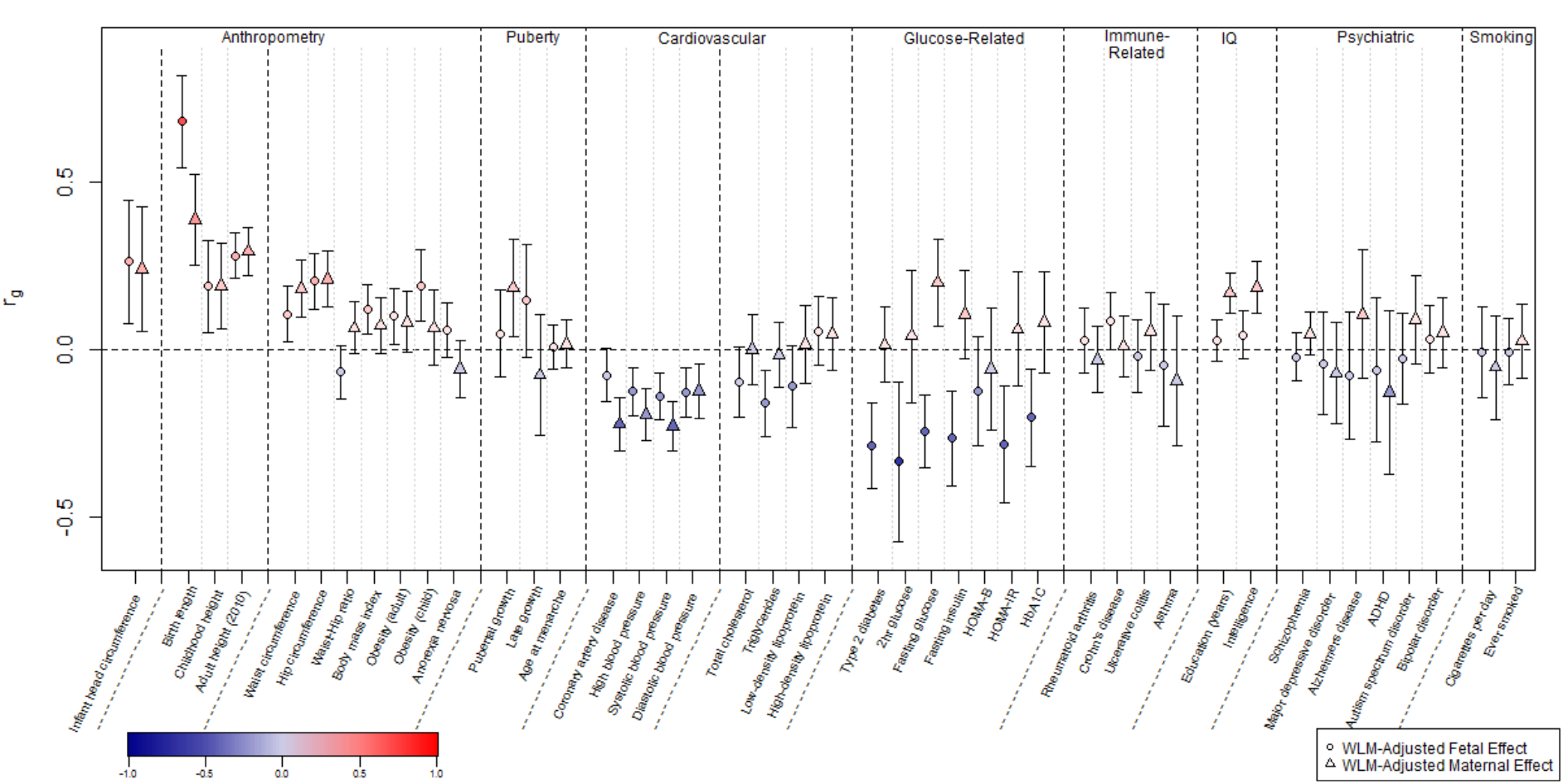
Genome-wide genetic correlation between birth weight (BW) and a range of traits and diseases in later life. Genetic correlation (r_g_) and corresponding 95% confidence intervals between BW and the traits were estimated using linkage disequilibrium (LD) score regression in LD Hub. Genetic correlations were estimated from the summary statistics of the weighted linear model (WLM)-adjusted fetal genome-wide association study (GWAS; WLM-adjusted fetal effect on own BW) and the WLM-adjusted maternal GWAS (WLM-adjusted maternal effect on offspring BW). The genetic correlation estimates are colour coded according to their intensity and direction (red for positive correlation and blue for negative correlation). HOMA-B/IR, homeostasis model assessment of beta-cell function/insulin resistance; HbA1c, hemoglobin A1c; ADHD, attention deficit hyperactivity disorder. See **Supplementary Table 10** for the references for each of the traits and diseases displayed and the genetic correlation results for other traits and diseases.

### Using genetics to estimate causal effects of intrauterine exposures on birth weight

The separation of direct fetal genotype effects from indirect maternal genotype effects on BW offers the novel opportunity to estimate the unconfounded causal influences of intrauterine exposures using Mendelian randomization (MR) analyses. The principle of MR is similar to that of a randomized controlled trial: parental alleles are randomly transmitted to offspring and are therefore generally free from confounding^22,23^. Consequently, an association between a maternal genetic variant for an exposure of interest, and offspring BW, after accounting for fetal genotype, provides evidence that the maternal exposure is causally related to offspring BW (**Figure 3A**). Previous attempts to estimate causal effects of maternal exposures on offspring BW were limited by an inability to adjust for fetal genotype in adequately-powered samples^24^. However, this limitation can now be overcome by using WLM-adjusted estimates in a two-sample setting. We applied two-sample MR^25^ to estimate causal effects of maternal exposures on offspring BW, focusing on height, glycemic traits and SBP. We selected SNPs known to be associated with each exposure, and regressed the WLM-adjusted maternal effect sizes on BW for those SNPs against the effect estimates for the maternal exposure, weighting by the inverse of the variance of the maternal exposure effect estimates. In the same way, we used the WLM-adjusted fetal effects to estimate the casual effect of the offspring’s genetic potential on their own BW, and compared the results with the estimated maternal causal effects.

**Figure 3:**
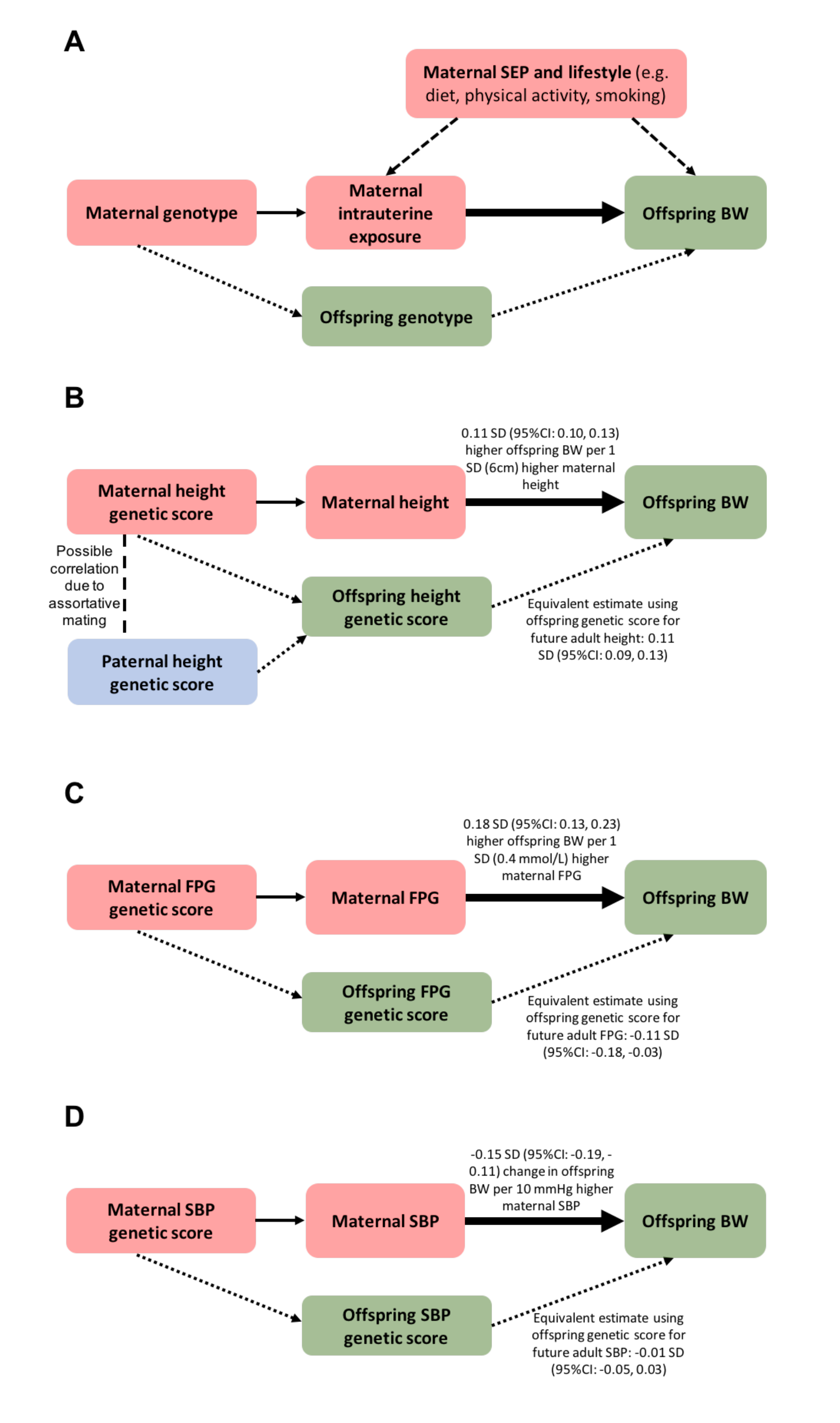
Mendelian randomization to assess the causal effect of maternal intrauterine exposures on offspring birth weight (adapted from Lawlor et al. ^44^) a. Diagram demonstrating how Mendelian randomization may be used to assesses the causal effect of a maternal exposure on offspring birth weight (BW). The analysis assumes that: (i) maternal genotype (instrumental variable, IV) is robustly associated with maternal (intrauterine) exposure; (ii) confounders of the maternal exposure-offspring outcome association are not associated with the maternal genetic IV; (iii) the maternal genetic IV is only associated with offspring BW through its association with the maternal intrauterine exposure, and through no other pathway. Since maternal and fetal genotypes are correlated (r = 0.5), it is essential to account for the offspring genotype in this analysis. The continuous, thin arrow represents the relationship between the genetic instrument and intrauterine exposure. The dashed arrows represent potential confounding via maternal characteristics. The dotted arrows represent potential violation of assumption 3 via the offspring genotype. The thick arrow represents the causal effect of interest estimated in Mendelian randomization analyses, after accounting for offspring genotype effects.
b. Higher offspring BW is caused both by direct fetal genetic effects of height-raising alleles and indirect effects of maternal height-raising alleles. Maternal indirect effects of height-raising alleles may increase offspring BW by increasing the space available for growth, but we cannot rule out alternative pathways e.g. the contribution of assortative mating, which would cause the maternal indirect association estimates to be correlated with direct fetal effects.
c. Higher maternal fasting glucose levels are causally associated with higher offspring BW. Conversely, in the offspring, a genetic score of alleles known to raise adult fasting glucose levels is associated with lower BW, indicating that these alleles have opposite maternal and fetal effects on BW. This is likely due to their effects on insulin: variants that lower maternal insulin levels lead to higher maternal glucose, which crosses the placenta and stimulates fetal insulin-mediated growth. However, the same variants in the fetus cause lower fetal insulin levels, and consequently, reduced fetal insulin-mediated growth.
d. Higher maternal blood pressure is causally associated with lower offspring BW. After adjusting for maternal effects, there was no evidence of an effect of the offspring’s own SBP genetic score on their own BW. SEP, socio-economic position; BW, birth weight; FPG, fasting plasma glucose; SBP, systolic blood pressure. 1 SD of BW = 484g^9,44^

### Height and birth weight

We used the WLM-adjusted estimates to investigate the relationship between maternal height and offspring BW. Classical animal experiments^26^ have demonstrated that larger maternal size can support greater fetal growth. This is supported by observational human data showing that offspring height shifts from being closer to maternal than paternal height percentile in infancy towards mid-parental height in adulthood, the latter reflecting the predominant role of inherited genetic variation^27^. However several observational studies have provided mixed evidence regarding correlations between maternal or paternal height and offspring BW: some studies show a stronger correlation with maternal than paternal height^28,29^, which would be consistent with a role for intrauterine effects (since both parents contribute equally to offspring genotype), while others show that maternal height is as strongly correlated with offspring BW as paternal height^30-32^. The MR analysis, using 693 height-associated SNPs as the instrumental variable^33^ (**Supplementary Table 14**), estimated that a 1 SD (6cm) higher maternal height is causally associated with a 0.11 SD (95%CI: 0.10, 0.13) higher offspring BW (**Figure 3B**), independent of the direct fetal effects. This estimate was similar in magnitude to that obtained using the WLM-adjusted fetal effects on own BW (0.11 SD (95%CI: 0.09, 0.13)), which reflects the role of inherited height alleles (**Supplementary Table 15**). Both a previous study^34^ and complementary analysis using transmitted and non-transmitted alleles in mother-offspring pairs (comparison of effects of maternal non-transmitted height alleles with alleles transmitted to offspring in N=3,485 and 4,962 mother-offspring pairs, respectively) estimated a much larger contribution of direct fetal effects than indirect maternal effects to offspring BW (**Supplementary Table 16**), however the sample sizes in both analyses were relatively small. To test whether the maternal height effect might be influencing BW by increasing gestational duration, as previously reported^34^, we applied the same MR analysis to maternal genotype effects on gestational duration^21^, but found little supportive evidence (P=0.12; **Supplementary Table 15**). The MR results from the current study are consistent with the hypothesis that greater maternal height causally increases BW, and that this effect is independent of the direct BW-raising effect of height alleles inherited by the fetus. For the maternal effect, we cannot rule out causal pathways other than the greater availability of space for fetal growth: causal associations between greater height and more favourable socio-economic position^35^, for example, could enhance maternal nutritional status and result in higher offspring BW. We also cannot exclude the contribution of assortative mating^36^ to these results: correlation between maternal and paternal height genotypes could lead to similar maternal and fetal MR estimates.

### Glycemic traits and birth weight

We used the WLM-adjusted estimates to assess the causal effect of maternal fasting glucose levels on BW with precision that was not achievable previously^24^ due to the inability to adjust for direct fetal effects in a large sample. Maternal glucose is a key determinant of fetal growth: it crosses the placenta, stimulating the production of fetal insulin which promotes growth^37^, and as a consequence, strong, positive associations are seen between maternal fasting glucose, or fetal insulin levels, and offspring BW^38^. In a randomized clinical trial of women with gestational diabetes mellitus, glucose control was shown to reduce offspring BW^39^. Therefore, we anticipated detecting a positive causal effect of maternal glucose on offspring BW. Indeed, the MR analysis using 33 fasting glucose-associated SNPs (**Supplementary Table 14**), estimated an 0.18 SD (95%CI: 0.13, 0.23) higher offspring BW due to 1 SD (0.4mmol/L) higher maternal fasting glucose, independent of the direct fetal effects (**Supplementary Table 15; Figure 3C**). A large part of the genetic variation underlying fasting glucose levels is implicated in pancreatic beta cell function and thus overlaps with the genetics of insulin secretion. To estimate the causal effect of insulin secretion on BW, we used 18 SNPs as instrumental variables that are associated with disposition index (DI), which is a measure of insulin response to glucose, adjusted for insulin sensitivity. Low values of DI are associated with higher T2D risk. Alleles that increase insulin secretion in the mother tend to decrease her glucose levels, which consequently reduces insulin-mediated growth of the fetus. This was reflected in the negative causal estimate from the MR analysis of the effect of maternal DI on offspring BW (−0.17 SD per 1 SD higher maternal DI (95%CI: - 0.26, −0.08); **Supplementary Table 15**). In contrast, we estimated that BW was 0.10 SD (95%CI: 0.02, 0.19) higher per 1 SD genetically higher fetal DI (**Methods**), highlighting that genetic variation underlying insulin secretion plays a key role in fetal growth, and suggesting that the genetic effects on DI are similar in fetal and adult life.

BW associations with previously-reported GWAS SNPs for fasting glucose, T2D, insulin secretion and insulin sensitivity loci were directionally consistent with the overall genetic correlations and supported the opposing contributions of fetal versus maternal glucose-raising alleles on BW (**Supplementary Figures 10-13**). Taken together with the WLM-adjusted genetic correlations, the MR results underline the importance of fetal insulin in fetal growth and demonstrate that fetal genetic effects link lower BW with reduced insulin secretion and higher T2D risk in later life^6^.However, further work will be needed to investigate the role of maternal indirect genetic effects in the relationship between high BW and higher future risk of T2D. The latter relationship may be driven by (i) maternal genetic predisposition to T2D resulting in raised glycemia in pregnancy and high offspring BW, then later offspring T2D through inheritance of maternal risk alleles, or (ii) a programming effect of exposure to high maternal glucose on later offspring T2D risk, or (iii) a combination of the two. The proportion of the negative BW-T2D covariance explained by fetal genotype effects on own BW was estimated to be 36% (95%CI: 15, 57; **Supplementary Table 17**), though this is likely an underestimate since current methods cannot adjust for the opposing effects of maternal genotypes.

### Blood pressure and birth weight

Observational studies of the relationship between BW and later life BP have produced mixed findings: some studies indicate that lower BW is associated with higher later-life BP and related comorbidities^40^, whereas others have shown that this relationship could be driven by a statistical artifact due to adjusting for current weight^41,42^. We previously showed that genetic factors account for a large proportion of an association between lower BW and higher BP^5^, but it was not clear whether this was due to direct fetal genotype effects, or indirect maternal effects, or a combination of the two. We explored this contested association further using several complementary analyses. The estimate of the BW-SBP covariance explained was higher when using the maternal genotyped SNP associations with offspring BW (65% (95%CI: 57, 74%)), than when using the fetal genotype associations with own BW (56% (95%CI: 48, 64%); **Supplementary Table 17**). A similar pattern was seen with the BW-DBP covariance (72% (95%CI: 58, 85%) explained using the maternal genotyped SNP associations with offspring BW and 56% (95%CI: 46, 67%) explained using the fetal genotype associations on own BW; **Supplementary Table 17**). Together with the larger maternal than fetal genetic correlations (**Figure 2**), these results point to the predominant importance of indirect maternal effects of BP genetics on offspring BW (**Supplementary Figures 14 and 15**). In line with this, MR analyses indicated that a 1SD (10mmHg) higher maternal SBP is causally associated with a 0.15 SD (95%CI: −0.19, −0.11) lower offspring BW, independent of the direct fetal effects. In contrast, there was no fetal effect of SBP on their own BW, after adjusting for the indirect maternal effect (−0.01 SD per 10mmHg, 95% CI: −0.05, 0.03; **Supplementary Tables 14 and 15; Figure 3D**). Similar results were seen in the WLM-adjusted MR analyses of DBP on both offspring and own BW.

### Estimating the causal effect of BW-lowering intrauterine exposures on offspring SBP

Having established (i) substantial negative genetic covariance between BW and SBP and (ii) indirect causal effects of maternal SBP-raising genotypes on lower offspring BW, a key question is whether maternal SNPs that reduce offspring BW through intrauterine effects are also associated with higher SBP in their adult offspring. Such an association would suggest that the maternal intrauterine effects also cause the later BP effect (i.e. possibly through developmental adaptations) (**Figure 4A; Supplementary Figure 16**). To investigate this possibility, we tested the conditional association between maternal and offspring genetic scores for BW and offspring SBP as measured in 3,886 mother-offspring pairs in the UK Biobank, with sensitivity analyses in 1,749 father-offspring pairs. The fetal genetic score for lower BW was associated with higher offspring SBP, even after adjustment for maternal (or paternal) BW genetic score. However, when adjusted for fetal genotypes, the maternal allele score for lower BW was associated with lower (*not higher*) offspring SBP (**Supplementary Table 18**). Taken together, our results demonstrate that the observed negative correlation between BW and later SBP is driven by (i) the causal effect of higher maternal SBP on lower offspring BW (**Figure 3D**), in combination with (ii) the subsequent transmission of SBP-associated alleles to offspring, which then increase offspring SBP (**Figure 4B**), rather than by long-term developmental compensations to adverse *in utero* conditions.

**Figure 4:**
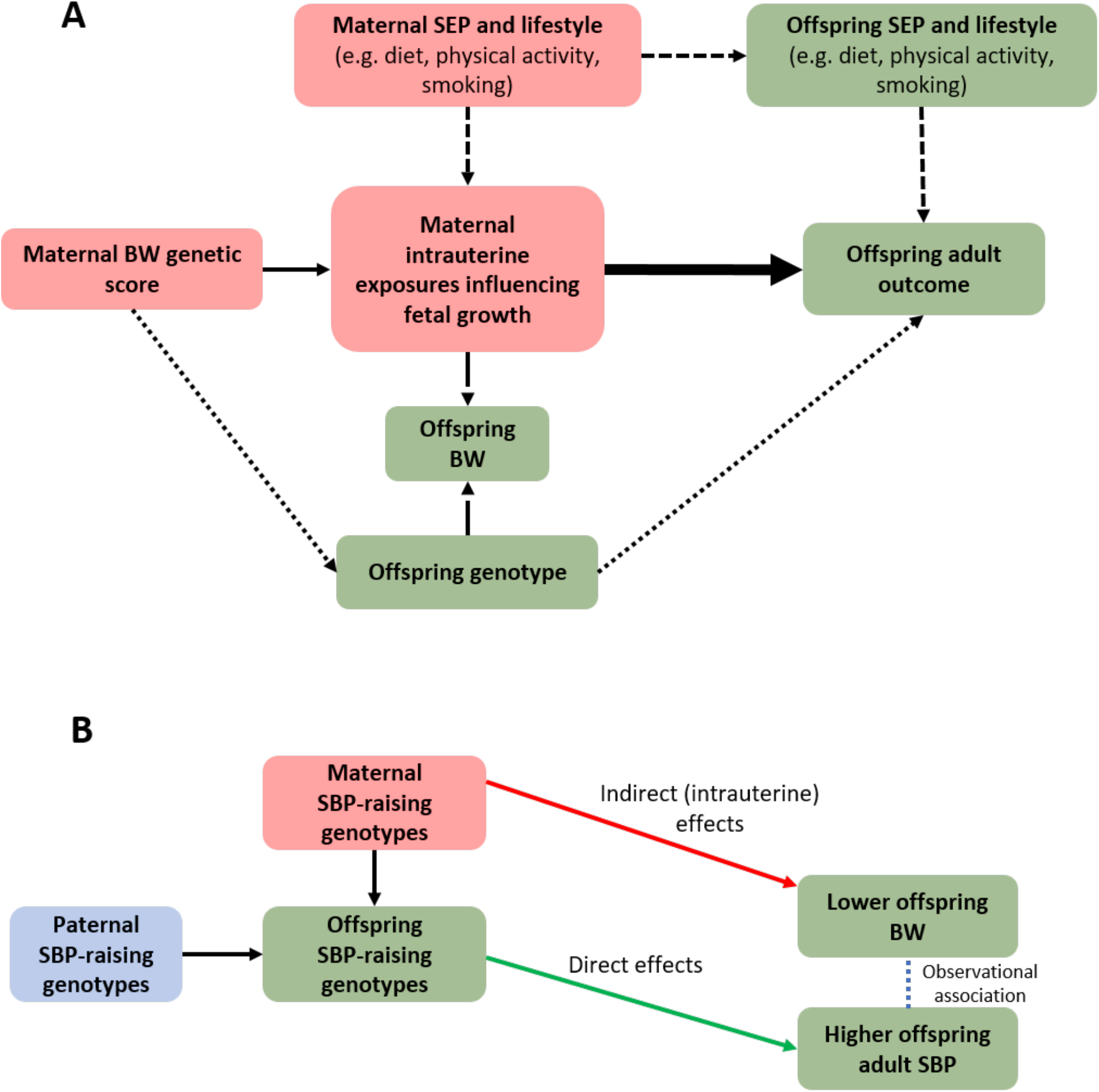
Mendelian randomization to assess the causal effect of intrauterine growth on offspring adult outcomes, using maternal intrauterine exposures that influence fetal growth. A. Diagram demonstrating how Mendelian randomization may be used to assesses the causal effect of intrauterine exposures that affect fetal growth on later-life offspring outcomes. The analysis assumptions are the same as in any Mendelian randomization analysis (see **Figure 3**). The instrumental variable should be associated with offspring birth weight (BW) independently of offspring genotype, so it is again essential to adjust the analysis for the offspring genotype. The continuous, thin, arrow represents the relationship between the genetic instrument and intrauterine exposure. The long-dashed arrows denote the (maternal and possibly fetal) genotype associations with BW; these arrows highlight the assumption that the genetic variation is influencing the offspring adult outcome via intrauterine growth pathways, not BW. The short-dashed arrows represent potential confounding via maternal and offspring characteristics. The dotted arrow represents potential violation of assumption 3 of Mendelian randomization analysis (see **Figure 3** legend) via the offspring genotype. The thick arrow represents the causal effect of interest estimated in Mendelian randomization analyses, after accounting for offspring genotype effects. We have not specifically conducted this Mendelian randomization analysis as we do not have effect estimates for the SNP-maternal intrauterine exposures influencing fetal growth. However, we have used the presence/absence and direction of association in 3,886 mother-offspring pairs from the UK Biobank to indicate whether the intrauterine environment was causing changes in adult SBP (see **Supplementary Table 18** for full results).
B. Taken together, our results demonstrate that the observed negative correlation between BW and later SBP may be driven by the causal effect of higher maternal SBP on lower offspring BW (red arrow), in combination with the subsequent transmission of SBP-associated alleles to offspring (green arrow), which then increase offspring SBP, rather than by long-term developmental compensations to adverse *in utero* conditions. SEP, socio-economic position; BW, birth weight; SBP, systolic blood pressure.

## DISCUSSION

In greatly-expanded GWAS and follow-up analyses of own and offspring BW, we have identified 214 novel association signals and have partitioned the genetic effects on BW into direct fetal and indirect maternal (intrauterine) effects, both for genome-wide significant SNPs, and for SNPs across the genome. Further analyses using these partitioned effects indicated both fetal-specific and maternal-specific mechanisms and tissues involved in the regulation of BW, and also mechanisms with directionally-opposing effects in the fetus and mother (e.g. insulin secretion, fasting glucose).

The variety of phenotypes associated with the identified BW SNPs, and pathways highlighted, illustrate that fetal growth is the result of many different processes^43^. We used the knowledge that subsets of BW-associated SNPs influence BP while others influence glycemic traits, or height, together with the WLM-adjusted estimates of maternal and fetal effects on BW to achieve a deeper level of insight into the relationships between BW and these adult traits. MR analyses using the WLM-adjusted estimates showed (i) evidence that both direct fetal and indirect maternal effects of height-raising genotypes contribute to higher offspring BW, (ii) that fetal, and not maternal, genotype effects explain the negative genetic correlation between BW and later T2D, and (iii) that the negative genetic correlation between BW and adult SBP is the result of both indirect SBP-raising effects of maternal genotypes reducing offspring BW, and direct effects of fetal genotypes on higher adult SBP. The resolution of maternal vs. fetal effects was higher in these MR analyses than has previously been achieved using analyses of available mother-child pairs^44^, due to greater statistical power. Recently, a number of studies have attempted to use MR methodology to investigate causal links between BW and later T2D^45-47^. However, such naïve MR analyses using two-sample approaches in unrelated sets of individuals, which do not properly account for the correlation between maternal and fetal genotype effects, may result in erroneous conclusions regarding causality. Future investigations into causal links between BW and later T2D or other disease outcomes will require larger samples than are currently available, that have maternal and offspring genotypes in addition to offspring later-life disease outcomes.

There are some limitations to this study. Although we were able to fit the full SEM at the 305 genome-wide significant SNPs, we were unable to fit the SEM at all SNPs across the genome. We have shown previously how a two degree of freedom test based on this SEM (i.e. where maternal and fetal paths are constrained to zero) can have greater power to detect associated loci, particularly when maternal and fetal genetic effects on the phenotype are similar in magnitude (including situations where the effects operate in opposite directions). However, we are currently unable to fit the SEM nor conduct an equivalent test in a computationally feasible manner across the genome. If such a test were developed, it would provide greater power than the current one degree of freedom tests used in the WLM-adjusted analyses, particularly for SNPs where maternal and fetal genetic effects operate in opposite directions, and could therefore be used for locus detection in future analyses. Additionally, there are a number of limitations relating to the MR analyses. First, the MR results concern BW variation within the normal range and do not necessarily reflect the effects of extreme environmental events (e.g. famine), which may exert qualitatively different effects and produce long-term developmental compensations in addition to low BW. Additionally, we have assumed a linear relationship between BW and later life traits, which is an oversimplification, particularly for T2D: higher BW is associated with later T2D risk, in addition to lower BW, particularly in populations with a high prevalence of T2D. MR is not well placed to examine the effects of extreme events, or non-linear relationships, and alternative methodology will be necessary to investigate life-course associations in this context. Second, BW is the end marker of a developmental process, with critical periods during the process that may make the fetus particularly sensitive to environmental influences. The MR analyses could therefore be masking effects at certain critical periods. We would need to look at maternal exposures on intrauterine growth trajectories or the specific function of the genetic variants on BW to interrogate this further. Third, we have assumed that genetic variants identified in large GWAS of SBP and glycemic traits in males and non-pregnant females are similarly associated in pregnant women. This assumption is reasonable, given that genetic associations are generally similar in pregnant vs non-pregnant women, though there is some indication that genetic effects on SBP are weaker in pregnancy (see Table 2, eTable and eTable 6f in Tyrrell et al. ^24^). Fourth, we have not investigated the potential gender difference in the associations between BW and later life traits. There is evidence that the association between BW and both T2D^48^ and SBP^49^ is stronger in females than males. However, to perform the MR analyses, we would require male and female-specific effect sizes for each of the exposures, which are currently not available. Finally, we have assumed that the critical period of exposure to maternal indirect genetic effects is pregnancy, and that the estimates do not reflect pre-pregnancy effects on primordial oocytes or post-natal effects^44^. However, since we have used BW-associated SNPs, the maternal effects are most-likely mediated *in utero*. While we cannot rule out postnatal effects^50^, our analysis of offspring SBP associations with BW-associated SNPs in father-child pairs showed different associations compared with mother-child pairs, implying postnatal effects were unlikely.

To conclude, the systematic separation of fetal from maternal genetic effects in a well-powered study has enhanced our understanding of the regulation of BW and of its links with later cardiometabolic health. In particular, we show that the association between lower BW and higher adult BP is attributable to genetic effects, and not to intrauterine programming. In successfully separating fetal from maternal genetic effects and using them in Mendelian randomization analyses, this work sets a precedent for future studies seeking to understand the causal role of the intrauterine environment in later-life health.

## ONLINE METHODS

### Ethics statement

All human research was approved by the relevant institutional review boards and conducted according to the Declaration of Helsinki. The UK Biobank has approval from the North West Multi-Centre Research Ethics Committee (MREC), which covers the UK. Participants of all studies provided written informed consent. Ethical approval for the ALSPAC study was obtained by the ALSPAC Ethics and Law Committee and the local Research Ethics Committees.

### Statistical tests

Details of statistical tests used in the various analyses are reported under the appropriate headings below. All tests were two-sided, unless otherwise stated.

### UK Biobank phenotype preparation

The UK Biobank is a study of 502,655 participants^51^. A total of 280,315 participants reported their own birth weight (BW) in kilograms at either the baseline visit or at least one of the follow-up visits. Participants reporting being part of a multiple birth were excluded from our analyses (N=7,706). For participants reporting BW at more than one visit (N=11,214), the mean value of the reported BWs was used, and if the mean difference between any 2 time points was >1kg, the participant was excluded (N=74). Data on gestational duration were not available; however, in order to exclude likely pre-term births, participants with BW values <2.5kg or >4.5kg were excluded (N=36,330). The remaining BW values were Z-score transformed separately in males and females for analysis.

Female participants were also asked to report the BW of their first child. A total of 216,839 women reported the BW of their first child on at least one assessment center visit. Values were recorded to the nearest whole pound, and were converted to kilograms for our analyses. Where women reported the BW of the first child at multiple time points (N=11,353) these were averaged and women were excluded if the mean difference between any 2 offspring BW measurements was >1kg (N=31). Women who reported the BW of their first child <2.2kg or >4.6kg were excluded (N=6,333). BW of first child was regressed against age at first birth and assessment center location. Residuals from the regression model were converted to Z-scores for analysis (sex of the first child was not available, so we were unable to calculate sex-specific Z-scores).

### UK Biobank ethnicity classification and genome-wide association analysis

We analysed data from the May 2017 release of imputed genetic data from the UK Biobank, a resource extensively described elsewhere^51^. Given the reported technical error with non-HRC imputed variants, we focused exclusively on the set of ~40M imputed variants from the HRC reference panel.

In addition to the quality control metrics performed centrally by the UK Biobank, we defined a subset of “white European” ancestry samples. To do this, we generated ancestry informative principal components (PCs) in the 1000 genomes samples. The UK Biobank samples were then projected into this PC space using the SNP loadings obtained from the principal components analysis using the 1000 genomes samples. The UK Biobank participants’ ancestry was classified using K-means clustering centered on the 3 main 1000 genomes populations (European, African, South Asian). Those clustering with the European cluster were classified as having European ancestry. The UK Biobank participants were asked to report their ethnic background. Only those reporting as either “British”, “Irish”, “White” or “Any other white background” were included in the clustering analysis. In total, 217,397 participants with a valid measure of their own BW and 190,406 women with a valid measure of BW of first child were classified as European and included in analyses. For trans-ethnic analyses all participants with valid phenotypes were included regardless of ancestry (N=227,530 participants with a valid measure of their own BW and N=210,208 with a valid measure of the BW of their first child).

Association analysis was conducted using a linear mixed model implemented in BOLT-LMM v2.3^52^ to account for population structure and relatedness. Only autosomal genetic variants which were common (MAF>1%), passed QC in all 106 batches and were present on both genotyping arrays were included in the genetic relationship matrix (GRM). For the genome-wide association study (GWAS) of the participants’ own BW, genotyping array and year of birth were included as covariates in all models. For the GWAS of the BW of the first child, genotyping array and genotyping release (interim vs. full) were included as covariates in the regression model, and indels, regions of long range LD (as defined in ^51^) and SNPs with Hardy-Weinberg equilibrium P-values<1×10^−6^ were excluded from the GRM.

### GWAS of own birth weight

#### European ancestry meta-analysis of own birth weight

The European ancestry GWAS meta-analysis of own BW consisted of two components: (i) 80,745 individuals from 35 studies participating in the EGG Consortium from Europe, USA and Australia; and (ii) 217,397 individuals of white European origin from the UK Biobank. Studies from the EGG Consortium conducted genome-wide association analysis of own BW that was Z-score transformed separately in males and females, and adjusted for study-specific covariates, including gestational duration, where available (**Supplementary Table 1**). GWASs were imputed up to the 1000 Genomes^53^ (1000G) reference panel. We combined the sex-specific BW association summary statistics across the EGG studies in a fixed-effects meta-analysis, implemented in GWAMA^54^, and subsequently combined the resulting summary statistics with the UK Biobank summary statistics using a second fixed-effects meta-analysis (max N=297,142). Variants failing GWAS quality control filters, reported in less than 50% of the total sample size in the EGG component, or with MAF<0.1%, were excluded from the European ancestry meta-analysis. We also performed a fixed-effects meta-analysis of the association summary statistics for 16,095 directly genotyped SNPs on the X-chromosome from the UK Biobank and the meta-analysis of the EGG studies (max N=270,929) using GWAMA^54^. A locus was defined as a gap of ≥500kb between any genome-wide significant SNPs, and the lead SNP within each locus was the SNP with the smallest P-value. The set of lead SNPs from each locus will be referred to as our genome-wide significant SNPs.

We were concerned that self-reported BW as adults in the UK Biobank would not be comparable with that obtained from more stringent collection methods used in the EGG studies. We conducted a heterogeneity test using Cochran’s *Q* statistic^55^, as implemented in GWAMA^54^, to assess the difference in allelic effects between the European EGG meta-analysis and the European subset of the UK Biobank, and we were unable to detect evidence of heterogeneity at lead SNPs after Bonferroni correction (all P>0.00029; **Supplementary Table 5**). However, we acknowledge that the power to detect evidence for heterogeneity using the Cochran’s *Q* statistic when comparing two groups is low and we use it here to highlight any SNPs with large differences in allelic effects. Although none of the SNPs reached the Bonferroni corrected threshold, there was an enrichment for low P, with more than double the expected number of SNPs with P<0.05 (18/173; **Supplementary Table 5**). In addition, the UK Biobank lacked information on gestational duration, which could impact the strength of association compared to the results obtained from the EGG studies which adjusted for gestational duration. Therefore, we conducted a further sensitivity analysis to specifically assess the impact of adjustment for gestational duration testing for heterogeneity in allelic effects at lead SNPs between EGG studies which adjusted for gestational duration (N=43,964) and the European subset of the UK Biobank. The only locus where the lead SNP showed significant heterogeneity, after Bonferroni correction, was rs1482852 at the *LOC339894/CCNL1* signal (*P_het_*=0.00015), which was a locus showing the strongest association with own BW and genome-wide significant in both EGG and the UK Biobank components independently.

There is potential for individuals to be in both the UK Biobank and EGG studies (i.e. the same individual in both the UK Biobank and a study within EGG) and this might lead to false positive association signals. We performed a bivariate linkage-disequilibrium (LD) score regression^19^ analysis using the European UK Biobank GWAS and European EGG meta-analysis summary statistics of own BW, and observed a regression intercept of 0.0266 (0.0077), indicating that the equivalent of approximately 3,524 individuals were in both GWAS analyses.

Univariate LD score regression^56^ of the European ancestry meta-analysis of own BW estimated the genomic inflation as 1.08, indicating that the majority of genome-wide inflation of the test statistics was due to polygenicity. To assess the impact of this inflation on the European ancestry meta-analysis, we re-calculated the association P-values after adjusting the test statistics for the LD score regression intercept. On the basis of this adjusted analysis, the lead SNP at 22 loci (out of 173) no longer reached genome-wide significance (**Supplementary Table 5**).

#### Approximate conditional and joint multiple-SNP (COJO) analysis to identify additional independent signals for own birth weight

Approximate COJO analysis^13^ was performed in GCTA^57^ using the European ancestry meta-analysis summary statistics to identify independent association signals attaining genome-wide significance (P<5×10^−8^). The LD reference panel was made up of 344,246 unrelated UK Biobank participants defined by the UK Biobank as having British ancestry and SNPs were restricted to those present in the HRC reference panel. At each locus, only SNPs labelled by GCTA as “independent” and not in LD with the original lead SNP (R^2^<0.05) were listed as secondary SNPs.

#### Trans-ethnic meta-analysis of own birth weight

To identify any further independent BW-associated SNPs, we conducted a trans-ethnic meta-analysis combining three components: (i) 80,745 individuals from the European ancestry component within EGG; (ii) 12,948 individuals from nine studies within EGG from diverse ancestry groups: African American, Afro-Caribbean, Mexican, Chinese, Thai, Filipino, Surinamese, Turkish and Moroccan; and (iii) 227,530 individuals of all ancestries from the UK Biobank. The same strategy and variant filtering criteria were applied as in the European meta-analysis of own BW (**Supplementary Figure 1**). None of the lead SNPs showed evidence of heterogeneity in BW allelic effects across the three components after Bonferroni correction (all Cochran’s *Q* P>0.169; **Supplementary Table 5**). Univariate LD score regression^56^ of the trans-ethnic meta-analysis estimated the genomic inflation as 1.08. After adjustment of the test statistics for the LD score regression intercept, the lead SNP at 8 loci (out of 11 that were added to our genome-wide significant loci from the trans-ethnic meta-analysis that were not identified in the European meta-analysis) dropped below genome-wide significance (**Supplementary Table 5**).

### GWAS of offspring birth weight

#### European ancestry meta-analysis of offspring birth weight

The European ancestry GWAS meta-analysis of offspring BW consisted of three components: (i) 12,319 individuals from 10 European GWAS imputed up to the HapMap 2 reference panel; and (ii) two European GWAS imputed up to the HRC panel (855 individuals from EFSOCH and 6,687 individuals from ALSPAC); and (iii) 190,406 individuals of white European origin from the UK Biobank. Studies from the EGG Consortium conducted genome-wide association analysis on offspring BW that was Z-score transformed, and adjusted for sex, gestational duration and ancestry informative principal components where necessary, (**Supplementary Table 3**). We then combined the BW association summary statistics across the 10 HapMap 2 imputed EGG studies in a fixed-effects meta-analysis, implemented in GWAMA^54^. We conducted a second European ancestry fixed-effects meta-analysis to combine the association summary statistics from the EGG meta-analysis with the UK Biobank, EFSOCH and ALSPAC (max N=210,267). The same strategy and variant filtering criteria were applied as in the meta-analysis of own BW. We also performed a fixed-effects meta-analysis of the association summary statistics for 18,137 directly genotyped SNPs on the X-chromosome from the UK Biobank and the meta-analysis of the EGG studies (max N=197,093) using GWAMA^54^. None of the lead SNPs showed evidence of heterogeneity in BW allelic effects, after Bonferroni correction (Cochran’s *Q* P>0.00060), between the UK Biobank and EGG studies and there was no enrichment for low P, with only 1/81 SNPs with P<0.05 (**Supplementary Table 5**). Using bivariate LD score regression^19^, we observed a regression intercept of 0.0165 (0.0063), indicating that the equivalent of approximately 1,015 individuals were in both the EGG and UK Biobank GWAS analyses of offspring BW.

Univariate LD score regression^56^ of the European ancestry meta-analysis estimated the genomic inflation as 1.05. Similar to the own BW GWAS results, we recalculated the P-values after adjusting the test statistics for this LD score intercept and the lead SNP at 8 loci (out of 81) dropped below genome-wide significance (**Supplementary Table 5**).

#### Approximate conditional and joint multiple-SNP (COJO) analysis to identify additional independent signals for offspring birth weight

We performed approximate COJO analysis^13^ using the European ancestry meta-analysis summary statistics of offspring BW, using the same reference panel as in the own BW analysis. Similarly to the analysis of own BW, SNPs labelled by GCTA as “independent” and not in LD with the original lead SNP (R^2^<0.05) were listed as secondary SNPs associated with offspring BW.

#### Trans-ethnic meta-analysis of offspring birth weight

We conducted a trans-ethnic meta-analysis combining three components: (i) 12,319 individuals from 10 European GWAS imputed up to the HapMap 2 reference panel; and (ii) two European GWAS imputed up to the HRC panel (855 individuals from EFSOCH and 6,686 individuals from ALSPAC); and (iii) 210,208 individuals of all ancestry from the UK Biobank. The same strategy and variant filtering criteria were applied as in the European meta-analysis of offspring BW (**Supplementary Figure 2**). None of the lead SNPs showed evidence of heterogeneity in BW allelic effects, after Bonferroni correction (Cochran’s *Q* P>0.00054), between the UK Biobank and EGG studies; however, there was an enrichment for low P, with 3/14 SNPs with P<0.05 (expected 1; **Supplementary Table 5**). Univariate LD score regression^56^ of the trans-ethnic meta-analysis estimated the genomic inflation as 1.04. We adjusted the test statistics for this LD score regression intercept, and the corresponding adjusted P-values for the lead SNP at 6 loci (out of 14 that were added to our genome-wide significant loci from the trans-ethnic meta-analysis that were not identified in the European meta-analysis) dropped below genome-wide significance (**Supplementary Table 5**).

### Colocalization methods

For each signal where we identified different lead SNPs in the GWAS of own BW and offspring BW, we performed co-localization analysis using the method implemented in the "coloc" R package^58^. For each signal, we input the regression coefficients, their variances and SNP minor allele frequencies for all SNPs 500kb up and downstream of the lead SNP from the European meta-analysis. We used the coloc.abf() function to calculate posterior probabilities that the own BW and offspring BW lead SNPs were independent (H_3_) or shared the same associated variant (H_4_). Default values were used for the prior probabilities in the coloc.abf() function. We call variants the same signal if the H_4_ posterior probability was greater than 0.50, and different signals if the H_3_ posterior probability was greater than 0.50.

### Estimation of genetic variance explained

Firstly, we estimated the proportion of BW variance explained by fetal genotypes, maternal genotypes and the covariance between the two at the 278 genome-wide significant signals in the Norwegian Mother and Child Cohort Study (MoBa-HARVEST; N=13,934 mother-offspring pairs; https://www.fhi.no/en/studies/moba/). This sample was independent of samples contributing to the discovery meta-analyses, apart from a small potential overlap with mothers from the MoBa-2008 sample that was included in the GWAS of offspring BW (affecting a maximum of 0.07% of the meta-analysis sample). For the 27 signals that had a maternal and fetal SNP, the fetal SNP was used in the analysis. This was to avoid any collinearity in the model due to the high correlation between the maternal and fetal SNPs. One SNP, rs77553582, was not available in MoBa-HARVEST, so we used a proxy SNP, rs2024344, in the analysis (r^2^=0.998 between rs77553582 and rs2024344). We excluded multiple births, babies of non-European descent, born before 37 weeks of gestation, born with a congenital anomaly or still-born. BW was Z-score transformed and all models were adjusted for sex, gestational duration and the first 4 ancestry informative principal components. We conducted a linear regression analysis in R^59^ using 13,934 mother-offspring pairs where offspring BW was regressed on the maternal and fetal genotypes at all 278 SNPs simultaneously. The proportion of variance explained by fetal genotypes at the 278 genome-wide significant signals was calculated as:

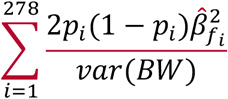

Where *p_i_* is the effect allele frequency of the i^th^ SNP, *β*̂_*fi*_ is the regression coefficient for the effect of the offspring’s genotype at the i^th^ SNP on offspring BW and *var(BW)* is the variance of offspring BW (which is approximately 1 as BW was Z-score transformed). A similar formula was used to calculate the variance explained by maternal genotypes:

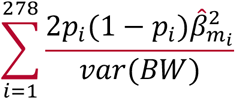

Where *β*̂_*mi*_ is the regression coefficient for the effect of the maternal genotype at the i^th^ SNP on offspring BW. Finally, a similar formula was used to calculate twice the covariance:

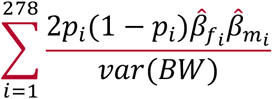

Secondly, we used maternal genome-wide complex trait analysis^11^ (M-GCTA) to estimate the proportion of variance explained in BW by genome-wide SNPs, or SNPs they tag, in the fetal genome, the maternal genome, the covariance between the two or environmental factors in MoBa-HARVEST. The same phenotype was used as in the previous analysis and the model was adjusted for sex and gestational duration. Mothers or offspring were excluded if they were related to others in the sample, using a genetic relationship cut-off 0.025, leaving N=7,910 mother-offspring pairs available for analysis.

### Identifying eQTL linked genes

To identify specific eQTL linked genes, we used the FUSION tool^60^ on the v6p release of the GTEx data^15^. FUSION is a gene-based data aggregation and integration method which incorporates information from gene-expression data and GWAS data to translate evidence of association with a phenotype from the SNP-level to the gene. Only gene level results from the adjusted model were taken forward for consideration. The threshold for statistical significance was estimated using the Bonferroni method for multiple testing correction across all tested tissues (tissue N=44, P<6×10^−7^). Each of the genes implicated by this analysis survived multiple test correction and were independent from other proximal genes tested in a joint model.

### Placenta eQTL look ups

We annotated genome-wide significant BW-associated SNPs with gene expression data (293/305 SNPs available) from placental samples of European ancestry from the Rhode Island Child Health Study^16^ (RICHS; N=123 with fetal genotype, including 71 with BW appropriate for gestational age, 15 small for gestational age, and 37 large for gestational age). We annotated genome-wide significant BW-associated SNPs on our list that had genome-wide empirical FDR<0.01 for association with one or more transcripts and r^2^>0.7 with a lead eQTL SNP.

### TAD pathways

Topologically associating domains (TAD) pathway analysis was performed using software described in Way *et al.*^17^. Briefly, the software uses publicly available TAD boundaries, identified in human embryonic stem cells and fibroblasts using a Hidden Markov Model^18^, to prioritize candidate genes at GWAS SNPs. These TAD boundaries are stable across different cell types and therefore can be used to identify genomic regions where non-coding causal variants will most likely impact tissue-independent function.

### Structural equation model for estimating adjusted maternal and fetal effects of the genome-wide significant variants

The structural equation modelling (SEM) approach used to estimate maternal and fetal effects that are independent of the fetal and maternal genotype respectively has been described elsewhere^12^. Briefly, to estimate the parameters for the SEM-adjusted fetal and maternal effects on BW, we use three observed variables available in the UK Biobank; the participant’s genotype, their own self-reported BW, and in the case of the UK Biobank women, the BW of their first child (**Supplementary Figure 17**). Additionally, the model comprises two latent (unobserved) variables, one for the genotype of the UK Biobank participant’s mother and one for the genotype of the participant’s offspring. From biometrical genetics theory, these latent genetic variables are correlated 0.5 with the participant’s own genotype, so we fix the path coefficients between the latent and observed genotypes to be 0.5. Participants who only report their own BW (including males), contribute directly to estimation of the fetal effect of genotype on BW and also indirectly to estimation of the maternal effect on BW since their observed genotype is correlated with their mother’s unmeasured latent genotype at the same locus. Similarly, summary statistics from the EGG meta-analysis of the unadjusted fetal effect (i.e. the European GWAS meta-analysis of own BW) can be incorporated into the model in this manner. Participants who report only their offspring’s BW (including mother’s reporting BW of their male offspring), contribute directly to estimation of the maternal effect on BW and indirectly to the estimate of the fetal effect on BW, since their observed genotype is correlated with their offspring’s latent genotype at the same locus. Again, summary statistics from the EGG meta-analysis of the unadjusted maternal effect (i.e. the European GWAS meta-analysis of offspring BW) can be incorporated into the model this way. These five components are fit to the five subsets of data (i.e. the UK Biobank participants with complete data, the UK Biobank participants with their own BW and genotype data only, EGG summary statistics for the unadjusted fetal effect of genotype on BW, the UK Biobank participants with their offspring’s BW and maternal genotype only and EGG summary statistics for the unadjusted maternal effect of genotype on BW) and then the likelihoods from each subset are combined. In addition to fitting the SEM to estimate the SEM-adjusted maternal and fetal effects, we fit a second model constraining the maternal and fetal effects to be zero and conducted a two degree of freedom Wald test to assess any effect of the SNP on BW. There is likely to be measurement error in the BW data in the UK Biobank, as well as some of the EGG studies, due to difficulty recalling BW. Additionally, the women in UK Biobank were asked to recall their offspring BW to the nearest pound. We have shown using simulations that both random measurement error (for example, due to difficulty in recall) and measurement error in offspring BW due to rounding to the nearest pound do not have a substantial influence on the estimation of either the maternal or fetal effects (see Warrington et al. ^12^). We therefore do not think that the imprecision of the UK Biobank BW data will substantially influence the results of downstream analyses.

The SEM was fit to data from 210 genome-wide significant fetal and 105 maternal SNPs from the GWAS meta-analysis; rs77553582, was only available in the GWAS of own BW from the EGG consortium so the SEM was not fit for this SNP (**Supplementary Figure 4**). In order to identify a subset of unrelated individuals in the UK Biobank (as the SEM cannot easily account for relatedness), we generated a genetic relationship matrix in the GCTA software package^57^ (version 1.90.2) and excluded one of every pair of related individuals with a genetic relationship greater than 9.375% (i.e. approximately half-way between third and fourth degree relatives). This gave us a subset of 382,001 unrelated individuals of European descent, after the same exclusions were made as in the GWAS, of which 85,518 individuals had reported their own BW and their offspring’s BW, 98,235 individuals reported their own BW only, and 73,981 reported their offspring’s BW only (the remaining 124,267 unrelated European individuals reported neither so were excluded from the analysis). We fit linear regression models to BW and offspring BW in the European, unrelated subset of individuals and adjusted for sex (own BW only), assessment centre and the top 40 ancestry informative principal components provided by the UK Biobank to account for any remaining population substructure. The residuals from this regression models were Z-score transformed for analysis. Because we included the summary statistics from the meta-analysis of the EGG studies, rather than the individual level data, we were unable to account for the small subset individuals who contributed to both the own BW and offspring BW GWAS meta-analyses. Based on the results from simulations (not shown), we expect that this non-independence will result in very slightly smaller standard errors and increased type 1 error rate, particularly for the fetal effect which is estimated from a larger sample size than was available to estimate the maternal effect. Therefore, we conducted a sensitivity analysis that first excluded EGG studies from the meta-analysis of own BW that contributed to both GWAS meta-analyses of own and offspring BW (e.g. ALSPAC), and then refitted the non-overlapping data in the SEM; these results are presented in **Supplementary Table 19**. For the four genome-wide significant SNPs identified on the X chromosome, we fit a slightly different SEM due to males having double the expected genetic variance at X linked loci compared to females. We did not incorporate summary statistics from the EGG consortium (since GWAS results were not stratified according to sex), so the model only includes the individual level data from the UK Biobank (additional details on the X chromosome analysis are provided in the **Supplementary Material** and **Supplementary Figure 18**).

We used the estimates from the SEM to classify SNPs into the following five categories; 1) ***fetal only***: the 95% confidence interval surrounding the fetal effect estimate does not overlap zero and does not overlap the 95% confidence interval around the maternal effect estimate. Additionally, the 95% confidence surrounding the maternal effect estimate overlaps zero; 2) ***maternal only***: the 95% confidence interval surrounding the maternal effect estimate does not overlap zero and does not overlap the 95% confidence interval around the fetal effect estimate. Additionally, the 95% confidence surrounding the fetal effect estimate overlaps zero; 3) ***fetal and maternal, effects going in the same direction***: the 95% confidence intervals around both the maternal and fetal effect estimates do not overlap zero, and their effect is in the same direction; 4) ***fetal and maternal, effects going in opposite direction***: the 95% confidence intervals around both the maternal and fetal effect estimates do not overlap zero, and their effects are in opposite directions; and 5) ***unclassified***: SNPs that do not fall into any of these categories, and therefore the 95% confidence intervals around the maternal and fetal effect estimates overlap, and at least one overlaps zero.

### Meta-analysis of maternal and fetal effects from a conditional regression analysis in mother-offspring pairs

We conducted conditional association analyses for all 305 genome-wide significant SNPs in 18,873 mother-offspring pairs from three studies (MoBa-HARVEST, ALSPAC and EFSOCH) adjusting for both maternal and offspring genotype and combined the summary statistics for each SNP in a fixed effects meta-analysis using METAL^61^. We compared the estimates of the maternal and fetal effects of this meta-analysis to the SEM-adjusted maternal and fetal effects using a heterogeneity test, and the results are presented in **Supplementary Table 6**.

### Approximation of the SEM for genome-wide analyses

The SEM is computationally intensive to fit, making it difficult to run on all SNPs across the genome. Therefore, we developed an approximation of the SEM using a linear transformation of the BW values and ordinary least squares linear regression, which we refer to as the weighted linear model adjusted (WLM-adjusted) analyses. The full details of the derivation are provided in the **Supplementary Material**. Briefly, from ordinary least squares regression we know that the estimated fetal effect size from the GWAS of own BW, 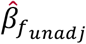, is calculated by dividing the sample covariance between BW and SNP by the sample variance of the SNP. Similarly, the estimated maternal effect from the GWAS of offspring BW, 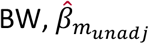, is calculated by dividing the sample covariance between offspring BW and SNP by the sample variance of the SNP. It follows that an estimate of the fetal effect adjusted for the maternal genotype is (see **Supplementary Material** for full derivation):

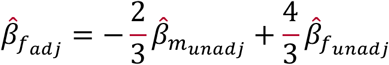

and an estimate of the maternal effect adjusted for the fetal genotype is:

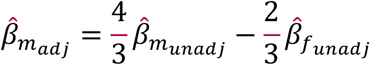

If the model is truly linear, then the same estimates can be obtained by transforming the reported BWs rather than the regression coefficients^62^. Similar to the SEM analyses, BW Z-scores in the UK Biobank participants were calculated from residuals of a regression model adjusting for sex (own BW only) and assessment centre, after the same exclusions were made as in the GWAS. For the UK Biobank participants who reported both their own BW and the BW of their offspring (N=101,541), we combined their BW Z-scores using the above formulae and conducted a GWAS in BOLT-LMM ^52^ to directly estimate the WLM-adjusted fetal and maternal effects for each SNP (see **Supplementary Figure 19** for a flow diagram of the full analysis pipeline). For the UK Biobank participants who only reported their own BW (N=115,070), we conducted a GWAS of their own BW Z-score in BOLT-LMM to estimate the unadjusted fetal effect for each SNP and then meta-analyzed the results with the unadjusted fetal effect estimates from the EGG consortium using a fixed-effects, inverse-variance weighted meta-analysis in METAL^61^. We followed the same procedure using participants who only reported their offspring’s BW in the UK Biobank (N=88,846) and meta-analyzed the unadjusted maternal effect estimates with those from the EGG consortium. The UK Biobank sample sizes used in this analysis are larger than those used in the SEM as the GWAS analyses are conducted in BOLT-LMM and can therefore account for the complex cryptic relationships between individuals. To get the WLM-adjusted maternal and fetal effect estimates, we combined the meta-analysis results of the unadjusted maternal and fetal effects for each SNP using the formulae above and their corresponding standard errors (see **Supplementary Material**). Finally, we conducted another fixed-effects, inverse-variance weighted meta-analysis to combine the WLM-adjusted maternal and fetal effect estimates from the UK Biobank participants with both BW measures and the combined WLM-adjusted effect estimates from the UK Biobank and EGG meta-analysis. A comparison of the results using this WLM method and the full SEM for the genome-wide significant SNPs is presented in **Supplementary Figure 8**.

### Gene expression integration

In order to identify which tissue types were most relevant to genes involved in BW, we applied LD score regression to specifically expressed genes (“LDSC-SEG”)^20^. We used the summary statistics from the GWAS meta-analysis of own and offspring BW and the WLM-adjusted meta-analyses, where the summary statistics from the WLM-adjusted meta-analyses were used to obtain tissue specific enrichments un-confounded by maternal and fetal genetic sharing. The method has been described previously^20^, but in brief it takes each tissue, ranking genes by a t-statistic for differential expression, using sex and age as covariates, and excluding all samples in related tissues. It then takes the top 10% of ranked genes, and makes a genome annotation including these genes (exons and introns) plus 100kb on either side. Finally, it uses stratified LD score regression to estimate the contribution of this annotation to per-SNP BW heritability, adjusting for all categories in the baseline model. We computed significance using a block jackknife over SNPs, and corrected for the number of tissues tested.

### Gene-set enrichment analysis (MAGENTA)

Pathway-based associations using summary statistics from the GWAS meta-analysis of own and offspring BW and WLM-adjusted meta-analysis for both the maternal and fetal effect were tested using MAGENTA^63^. Briefly, the software maps each gene to the SNP with the lowest P-value within a 110kb upstream and 40kb downstream window. This P-value is corrected for factors such as SNP density and gene size using a regression model. Genes within the HLA region were excluded. The observed number of gene scores within a given pathway with a ranked gene score above a given threshold (95^th^ or 75^th^ percentile) was calculated. This statistic was compared with 1,000,000 randomly permuted pathways of the same size to calculate an empirical P-value for each pathway. We considered pathways with false discovery rate (FDR) < 0.05 to be of interest. The 3,230 biological pathways tested were from the BIOCARTA, Gene Ontology, KEGG, PANTHER and READTOME databases along with a small number of custom pathways.

### Gestational duration associations

We extracted the 305 genome-wide significant BW-associated SNPs from the summary statistics in a recent GWAS of gestational duration^21^. The full table of 23andMe summary statistics was obtained directly from 23andMe. For BW SNPs that were also associated with gestational duration (P<1.6×10^−4^, corrected for 305 tests), we followed them up in 13,206 mother-child pairs from the MoBa-HARVEST, ALSPAC and EFSOCH studies. Preterm births (gestational duration <37 weeks) were removed before analysis, and gestational duration and BW were both z-score transformed. We conducted linear regression analyses to test the association between maternal or fetal genotype (unadjusted genotype effects) and gestational duration, BW or gestational duration adjusted for BW. Additionally, we conducted linear regression analyses for the same three outcomes including both the maternal and fetal genotypes (adjusted effects). The association analysis results were combined using inverse variance weighted meta-analysis. We also combined the unadjusted maternal SNP-gestational duration associations with the 23andMe summary statistics from Zhang *et al*.^21^ using P value based meta-analysis implemented in METAL^61^.

### Association between birth weight-associated SNPs and a variety of traits in the UK Biobank

We performed GWAS on 78 traits in the UK Biobank using BOLT-LMM in an analogous way to analysis of own BW. Association statistics for the 305 genome-wide significant BW-associated SNPs were then extracted from the results. Phenotype definitions for the 78 traits are described in Frayling *et al*.^64^. Additionally, for the 305 genome-wide significant BW-associated SNPs, and those in high LD with the 305 SNPs (r^2^>0.8), we searched the NHGRI GWAS catalog (https://www.ebi.ac.uk/gwas/; accessed 16th January 2018) for reported GWAS associations with other traits. These are reported in **Supplementary Table 13**.

### Linkage-Disequilibrium (LD) score regression

LD score regression was used to estimate the genetic correlation between two traits/diseases and has been described in detail elsewhere^19^. Briefly, the LD score is a measure of how much genetic variation each variant tags; so if a variant has a high LD score then it is correlated with many nearby variants. Variants with high LD scores are more likely to tag true signals and hence provide greater chance of overlap with genuine signals between GWAS. The method uses summary statistics from the GWAS meta-analyses of BW and the other traits of interest, calculates the cross-product of test statistics at each SNP, and then regresses the cross-product on the LD Score. Bulik-Sullivan *et al.* ^19^ show that the slope of the regression is a function of the genetic correlation between traits. Individuals contributing to the summary statistics of both GWAS meta-analyses, and population stratification within either GWAS, will only influence the intercept of the regression and therefore not bias the genetic correlation.

We used LDHub^65^ (ldsc.broadinstitute.org/) to perform LD score regression between BW and a large range of traits and diseases. LDHub is a centralized database which contains curated summary statistics from GWAS analyses of over 775 traits and diseases, including the recent release of summary statistics from GWAS analyses of many phenotypes in the UK Biobank (http://www.nealelab.is/blog/2017/7/19/rapid-gwas-of-thousands-of-phenotypes-for-337000-samples-in-the-uk-biobank). Due to the different LD structure across ancestry groups, the summary statistics from the European only BW analyses were uploaded to LDHub and genetic correlations were calculated for all available phenotypes. We conducted four separate analyses in LDHub: one for the GWAS of own BW, one for the GWAS of offspring BW, one for the WLM-adjusted fetal effect on own BW and the final one for the WLM-adjusted maternal effect on offspring BW.

To calculate the genetic correlation between the maternal and fetal effect estimates from the unadjusted and WLM-adjusted analyses, and also between gestational duration and the WLM-adjusted maternal and fetal effects, we used the scripts provided by the developer (https://github.com/bulik/ldsc).

### Mendelian randomization analyses of maternal and fetal exposures on offspring birth weight

Two sample Mendelian randomization analyses were performed for a number of exposures with own BW or offspring BW as outcomes. The exposures included height (SD units, where 1SD = 6cm), fasting glucose (SD units, where 1 SD = 0.4mmol/L), disposition index of insulin secretion (calculated from oral glucose tolerance test (OGTT) results as Corrected Insulin Response x 10,000 / √ (Fasting Plasma Glucose x Fasting Insulin x Mean Glucose during OGTT x Mean Insulin During OGTT)^66^), insulin sensitivity (calculated as fasting insulin adjusted for BMI) and systolic and diastolic blood pressure (mmHg). The SNP-exposure associations were taken from external studies (**Supplementary Table 14**). The SNP-outcome associations were taken from the current European GWAS meta-analyses of own BW, offspring BW, WLM-adjusted fetal effect on own BW and WLM-adjusted maternal effect on offspring BW. Two sample Mendelian randomization regresses effect sizes of SNP-outcome associations against effect sizes of SNP-exposure associations, with an inverse-variance weighted (IVW) analysis giving similar results to the commonly used two-stage least squares analysis in a single sample^67^. We performed several sensitivity analyses to assess the impact of genetic pleiotropy on the causal estimates including MR-Egger^68^, Weighted Median (WM)^69^ and Penalized Weighted Median (PWM)^69^ approaches (see **Supplementary Table 15** for results). Details of the R code for the MR analyses are provided elsewhere^68,69^. Due to the strong negative correlation between estimates of the maternal and fetal genetic effects on BW, we conducted simulations to confirm that this correlation did not bias the results of downstream MR analyses; these simulations are described in the **Supplementary Material**.

### Transmitted/non-transmitted allele scores in ALSPAC

Allelic transmission was determined for 4,962 mother/offspring pairs in ALSPAC. We first converted maternal and fetal genotypes into best guess genotypes where SNPs of interest had been imputed. Where one or both of the mother/offspring pair were homozygous, allelic transmission is trivial to determine. Where both mother and offspring were heterozygous for the SNP of interest we used phase imputation generated using SHAPEIT2^70^ to examine the haplotypes in the region of the SNP of interest to determine allelic transmission. Weighted allele scores were then generated for maternal non-transmitted, shared (maternal transmitted) and paternally inherited fetal alleles for systolic blood pressure (SBP), diastolic blood pressure (DBP), fasting glucose, insulin secretion and insulin sensitivity. Associations were tested between the weighted allele scores and BW.

### Covariance between BW and adult traits explained by genotyped SNPs

The genetic and residual covariance between BW and several quantitative and/or disease phenotypes was calculated in the UK Biobank in the BOLT-LMM^52^ implementation of the REML method, using directly genotyped SNPs. We included 215,444 individuals with data on own BW and 190,406 with data on offspring BW. All individuals were classified as being of European ancestry. SNPs with minor allele frequency < 1%, evidence of deviation from Hardy-Weinberg equilibrium (P≤1×10^−6^) or overall missing rate > 0.015 were excluded, resulting in 524,307 SNPs for analysis. Ninety-five per cent confidence intervals for the proportion of covariance explained by variants directly genotyped were calculated as gcov/(gcov+rcov) ± 1.96*gcovSE/abs(gcov+rcov) where gcov is genetic covariance, rcov is residual covariance and gcovSE is the standard error for gcov and abs is the absolute value. Details of the phenotype preparation for the adult traits is provided in the **Supplementary Material**.

### Testing for association between maternal SNPs associated with offspring birth weight and later-life offspring blood pressure

Using the UK Biobank, we tested whether maternal SNPs associated with offspring BW were also associated with offspring blood pressure in later life. The UK Biobank released kinship information generated in KING^71^, which included the kinship coefficients and IBS0 estimates. We defined parent/offspring pairs using the kinship coefficient and IBS0 cut-offs recommended in Manichaikul *et al*.^71^. There were 6,276 parent/offspring pairs, of which 5,901 were of European descent and 5,635 unique pairs with blood pressure data (for parents who had multiple offspring with blood pressure data, only the oldest offspring was included in the analysis); 3,886 mother/offspring pairs and 1,749 father/offspring pairs. We tested the relationship between unweighted allelic scores of BW-associated SNPs in mothers and offspring SBP (see **Supplementary Material** for SBP phenotype preparation) before and after adjusting for offspring genotypes at the same loci. We examined unweighted allelic scores consisting of all autosomal genome-wide significant BW-associated SNPs available in the UK Biobank (300 SNPs), 103 autosomal SNPs that showed evidence of a maternal effect, and a subset of 44 autosomal SNPs that showed evidence only of maternal effects on BW. We also looked at the SNPs previously associated with SBP (**Supplementary Table 14**) as a sensitivity analysis to rule out the possibility of postnatal pleiotropic effects of SNPs contaminating our results, we also tested the relationship between allelic scores of BW-associated SNPs in fathers and offspring SBP after adjusting for offspring genotype. All analyses were adjusted for offspring age at SBP measurement, sex and assessment center.

## Data availability

The genotype and phenotype data are available upon application from the UK Biobank (http://www.ukbiobank.ac.uk/). Individual cohorts participating in the EGG consortium should be contacted directly as each cohort has different data access policies. GWAS summary statistics from this study are available on publication via the EGG website (https://egg-consortium.org/).

## Acknowledgements

Full acknowledgements and supporting grant details can be found in the Supplementary material.

## Author contributions

### Central analysis and writing team

N.M.W., R.N.B., M.H., F.R.D., K.K.O., M.I.M., J.R.P., D.M.E., R.M.F.

### Statistical analysis

N.M.W., R.N.B., M.H., F.R.D., Ø.H., C.Laurin, J.B., S.P., K.H., B.F., A.R.W., A. Mahajan, J.T., N.R.R., N.W.R., Z.Q., G-H.Ø.M., M.Vaudel, M.N., T.M.S., M.H.Z., J.P.B., N.G., M.N.K., R.L-G., F.G., T.S.A., L.P., R.R., V.H., J-J.H., L-P.L., A.C., S.M., D.L.C., Y.W., E.T., C.A.W., C.T.H., N.V-T., P.K.J., J.N.P., I.N., R.M., N.P., E.M.v.L., R.J., V.L., R.C.R., A.E., S.J.B., W.A., J.A.M., K.L.L., C.A., G.Z., L.J.M., J.Heikkinen, A.H.v.K., K.J.G., N.R.v.Z., C.M-G., Z.K., S.D., H.M., E.V.R.A., M.Murcia, S.B-G., D.M.H., J.M.Mercader, K.E.S., P.A.L., S.E.M., B.M.S., J-F.C., K.Panoutsopoulou, F.S., D.T., I.P., M.A.T., H.Y., K.S.R., S.E.J., P-R.L., A.Murray, M.N.W., E.Z., G.V.D., Y-Y.T., M.G.H., K.L.M., J.F.F., D.M.S., N.J.T., A.P.M., D.A.L., J.R.P., D.M.E., R.M.F.

### Genotyping

F.R.D., Ø.H., T.M.S., M.H.Z., N.G., R.L-G., L.P., J-J.H., L-P.L., J.W.H., X.E., L.M., L.B., C.S.M., C.Langenberg, J’a.L., R.A.S., J.H.Z., G.H., S.M.R., A.J.B., J.F-T., C.M-G., H.G.d.H., F.R.R., Z.K., P.M-V., H.M., E.V.R.A., M.Bustamante, M.A., P.K., M.Stumvoll, T.A.L., C.M.v.D., A.K., E.Z., S-M.S., G.W.M., H.C., J.F.W., T.G.V., C.E.P., E.E.W., T.D.S., T.L., P.V., H.B., K.B., J.C.M., F.R., J.F.F., T.H., O.P., A.G.U., M-R.J., W.L.L.Jr, G.D.S., N.J.T., N.J.W., H.H., S.F.G., T.M.F., D.A.L., P.R.N., K.K.O., M.I.M., J.R.P., D.M.E., R.M.F.

### Sample collection and phenotyping

F.R.D., B.F., C.J.M., J.C., J.P.B., M.N.K., R.L-G., F.G., R.R., I.N., H.M.I., J.W.H., L.S-M., C.R., B.H., M.Kogevinas, L.C., M-F.H., C.S.M., F.D.M., C.Langenberg, J’a.L., R.A.S., J.H.Z., S.M.R., C.M-G., H.G.d.H., Z.K., P.M-V., S.D., G.W., M.M-N., M.Standl, C.E.F., C.T., C.E.v.B., M.Bustamante, D.M.H., A.L., B.A.K., M.Bartels, J.S., R.K.V., S.M.W., B.L.C., A.T., K.F.M., A-M.E., T.A.L., A.K., H.N., K.Pahkala, O.T.R., B.J., G.V.D., S-M.S., G.W.M., J.F.W., T.G.V., M.Vrijheid, J-C.H., L.J.B., C.E.P., L.S.A., J.B.B., J.G.E., E.E.W., A.T.H., T.D.S., M.Kähönen, J.S.V., T.L., P.V., H.B., K.B., M.Melbye, E.A.N., D.O.M-K., J.F.F., V.W.J., C.Pisinger, A.A.V., M-R.J., C.Power, E.H., W.L.L.Jr, G.D.S., N.J.W., H.H., S.F.G., D.A.L., K.K.O., M.I.M., J.R.P.

### Study Design and Principal Investigators

J.P.B., I.N., H.M.I., L.S-M., X.E., B.H., J.M.Murabito, M.Kogevinas, L.C., M-F.H., F.D.M., M.A., A.T., M.Stumvoll, K.F.M., A-M.E., T.A.L., C.M.v.D., W.K., A.K., H.N., K.Pahkala, O.T.R., B.J., G.V.D., Y-Y.T., S-M.S., G.W.M., H.C., J.F.W., T.G.V., M.Vrijheid, E.J.d.G., H.N.K., J-C.H., L.J.B., C.E.P., J.Heinrich, L.S.A., J.B.B., K.L.M., J.G.E., E.E.W., A.T.H., T.D.S., M.Kähönen, J.S.V., T.L., D.I.B., S.S., P.V., H.B., K.B., J.C.M., M.Melbye, E.A.N., D.O.M-K., F.R., J.F.F., V.W.J., T.H., C. Pisinger, A.A.V., O.P., A.G.U., M-R.J., C. Power, E.H., W.L.L.Jr, N.J.T., A.P.M., N.J.W., H.H., S.F.G., T.M.F., D.A.L., P.R.N., S.J., K.K.O., M.I.M., J.R.P., R.M.F.

## Competing interests statement

A.A.V. is an employee of AstraZeneca. D.A.L. has received support from Medtronic LTD and Roche Diagnostics for biomarker research that is not related to the study presented in this paper. M.I.M. serves on advisory panels for Pfizer, NovoNordisk, Zoe Global; has received honoraria from Merck, Pfizer, NovoNordisk and Eli Lilly; has stock options in Zoe Global; has received research funding from Abbvie, Astra Zeneca, Boehringer Ingelheim, Eli Lilly, Janssen, Merck, NovoNordisk, Pfizer, Roche, Sanofi Aventis, Servier, Takeda.

